# CherryML: Scalable Maximum Likelihood Estimation of Phylogenetic Models

**DOI:** 10.1101/2022.12.21.521328

**Authors:** Sebastian Prillo, Yun Deng, Pierre Boyeau, Xingyu Li, Po-Yen Chen, Yun S. Song

## Abstract

Phylogenetic models of molecular evolution are central to diverse problems in biology, but maximum likelihood estimation of model parameters is a computationally expensive task, in some cases prohibitively so. To address this challenge, we here introduce CherryML, a broadly applicable method that achieves several orders of magnitude speedup. We demonstrate its utility by applying it to estimate a general 400 × 400 rate matrix for amino acid co-evolution at protein contact sites.

---

Phylogenetic models of molecular evolution have a plethora of applications in biology. For example, models of amino acid substitution enable the estimation of gene trees and protein alignments, among other important applications [1–9]. These models posit that molecules evolve down a phylogenetic tree according to a continuous-time Markov process parameterized by one or more rate matrices. These models differ in what kind of molecular data they describe (e.g., DNA, RNA, amino acid, or codon sequences), the use of site rate variation (such as invariable sites model, the Γ model, or the probability-distribution-free model [10, 11]), the treatment of indels [12], and the assumption of i.i.d. (independent and identically distributed) sites [13]. A common aspect of all these models is that maximum likelihood estimation (MLE) of model parameters is computationally expensive, in some cases prohibitively so.

The main computational challenge arises from the unobserved ancestral states. A typical step during MLE involves estimating the rate matrix *Q* given a set of *m* multiple sequence alignments (MSAs) and associated phylogenetic trees, and computing the log-likelihood alone requires marginalizing out the unobserved ancestral states, which is done with Felsenstein’s pruning algorithm [14]. If *s* denotes the state space size (e.g., *s* =20 for amino acids), *l* the typical sequence length, and *n* the typical number of sequences per MSA, then computing the likelihood with Felsenstein’s pruning algorithm has a cost of Ω(*mn* (*ls*^2^ + *s*^3^)) for the typical model. This comes from having to compute, for each of the Ω(*mn*) edges, (at least) one matrix exponential of cost Ω(*s*^3^), and a recursion step of cost Ω(*s*^2^) per alignment site.

MLE for these models is usually performed with either zeroth-order optimization [9] or with the Expectation-Maximization (EM) algorithm [15, 16], both of which require Felsenstein’s pruning algorithm. As a consequence, full MLE is typically slow and rarely performed in practice [9]. Instead, researchers choose to utilize generic rate matrices such as the popular LG matrix [5] – which was estimated more than a decade ago – to perform their phylogenetic analyses. For more complex models, estimation with current approaches is just infeasible. For example, learning a general 400 × 400 rate matrix describing the co-evolution of contacting residues in a protein is out of reach with current approaches.

To overcome these challenges, we here propose CherryML, a broadly applicable method to scale up MLE for general phylogenetic models of molecular evolution. CherryML hinges on two key ideas: *composite* likelihood and time *quantization*. We describe these ideas in turn below and illustrate them in Figure 1a.

**Figure 1:**
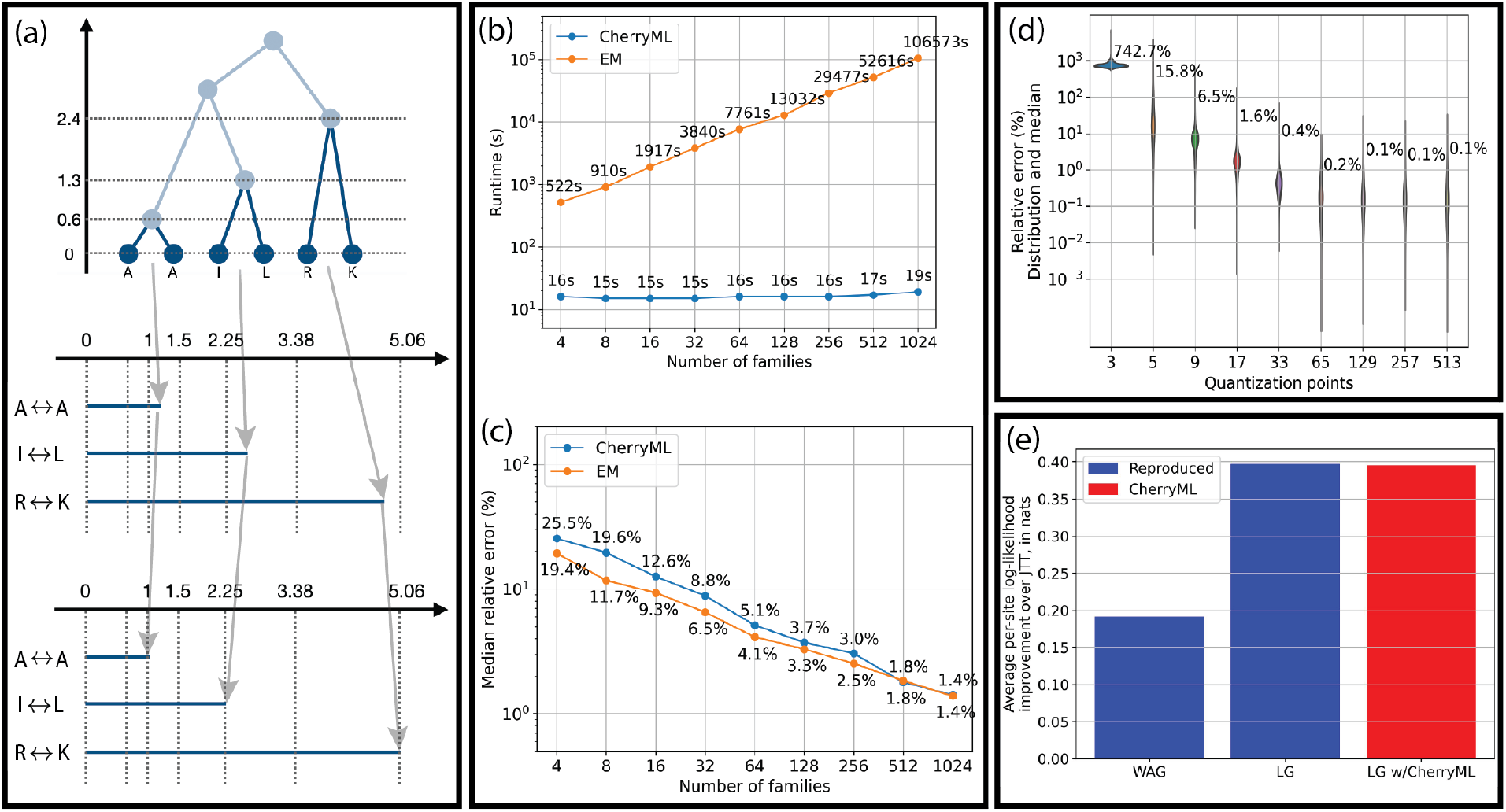
CherryML method applied to the LG model. (a) Sketch of the CherryML method as applied to a reversible model: transitions between cherries are treated as independent observations and quantized; here the quantization grid is {1, 1.5, 2.25, 3.38, 5.06 }. (b, c) Runtime and median estimation error as a function of sample size for our CherryML optimizer and the classical EM optimizer. The empirical loss of statistical efficiency for CherryML is small (<50%) while being more than a thousand times faster when applied to 1,024 families. Each family has 128 sequences. (d) On a large simulated dataset, time quantization error becomes negligible with as few as ≈ 100 quantization points. Distribution of relative error and median shown. (e) Using the evaluation protocol of the LG paper [5], we verified that the CherryML method produces comparable likelihoods on held-out families.

The first key idea underlying CherryML is to use a composite likelihood over cherries instead of the full likelihood. This means selecting cherries for each tree and replacing the full log-likelihood with the sum of log-likelihoods of the cherries (see Methods for full details). This reduces likelihood computation to *pairs* of sequences at a time, which is much simpler than the full likelihood. In particular, for a time-reversible model, the likelihood of a pair of sequences only depends on the distance between the nodes in the tree, and does not require marginalizing out ancestral states. Maximum composite likelihood estimation (MCLE) enjoys many of the properties of MLE, such as consistency under weak conditions [17].

The second key idea of CherryML is to quantize time. This means approximating transition times by one of finitely many values *τ*_1_ < *τ*_2_ < ⋯ < *τ*_*b*_. We choose the *τ*_*k*_ to be geometrically spaced and to cover the whole operational range of transition times; thanks to the geometric spacing, a few hundred values are enough to achieve a quantization error as low as 1% for all transition times. When time is quantized, the terms in the composite likelihood typically group together by quantized transition time *τ*_*k*_, and the cost of evaluating the likelihood no longer depends on the dataset size, dramatically speeding up optimization.

We applied the CherryML method to the seminal model of Le and Gascuel (LG) [5], which assumes that each site in every protein evolves under the same rate matrix *Q* and with a site specific rate. As a result, the composite log-likelihood has the functional form 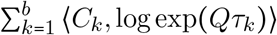 where *C*_*k*_ is the frequency matrix for transitions between states occurring at a quantized time of *τ*_*k*_. Hence, the cost of evaluating the likelihood no longer depends on the dataset size (see Methods for full details). This objective function can be easily optimized in modern first-order numerical optimization libraries such as PyTorch [18]. If *g* denotes the number of iterates of the first-order optimizer, the computational complexity of the CherryML method applied to the LG model is then Θ(*mnl* log *b* + *gbs*^3^). Here Θ(*mnl* log*b*) comes from computing the count matrices *C*_*k*_, and Θ(*gbs*^3^) comes from the first-order optimizer; the Θ(*mnl* log*b*) term will dominate for large datasets. In contrast, the cost of MLE with traditional methods such as zeroth-order optimization or EM is Ω(*gmn*(*ls*^2^ + *s*^3^)), and the Ω(*gmnls*^2^) term will dominate. Thus, ignoring constants, a back-of-the-envelope calculation reveals that CherryML will be at least 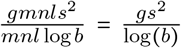 times faster than traditional methods, which is typically a massive speedup. Indeed, when learning a single-site model using *g* = 100 iterates and *b* = 100 quantization points for CherryML, the speedup is at least (up to constants) 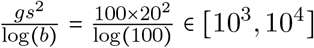 fold. When learning a co-evolution model, the speedup is at least (up to constants) 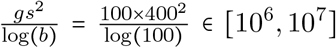 fold. The computational bottleneck of CherryML is computing the count matrices *C*_*k*_, and we have developed an efficient distributed-memory C + + implementation using MPI. (See Methods for the details.)

By using simulated data, we benchmarked the computational runtime and statistical efficiency of the CherryML method for the aforementioned LG model, and compared it with the performance of EM, for which we used the implementation in the Historian package [19]. The results are summarized in Figure 1b,c, which shows that CherryML is more than a thousand times faster than EM when run on 1,024 families with 128 sequences each, taking 19 seconds as compared to almost 30 hours. This comes only at the cost of less than 50% loss of statistical efficiency. Using data simulated on all 15,051 protein families studied in Yang *et al*. [20], we found that *b* ≈ 100 quantization points were enough to make the error introduced by time quantization negligible, as shown in Figure 1d.

We next applied CherryML to the Pfam data from Le and Gascuel [5]. We implemented their end-to-end estimation procedure, replacing the EM optimizer with the CherryML optimizer. We estimated phylogenetic trees and site-specific rates using FastTree [21], and then applied CherryML to estimate the 20 × 20 rate matrix *Q*_20_. Tree estimation and rate matrix estimation were iterated 3 times until convergence as typical, starting from the uninformative uniform rate matrix. We reproduced and extended the results from Figure 4 of Le and Gascuel [5] by adding our estimate of the LG rate matrix using our CherryML method. Figure 1e shows that the results obtained with the CherryML method are comparable to that obtained using EM.

Finally, we applied the CherryML method to estimate a general reversible 400 × 400 rate matrix *Q*_400_ describing the co-evolution of contacting sites in a protein. The mathematics of the model are similar to the LG model except that there is no site rate variation and *s* = 400 (see Methods for full details). The large size of the state space means that the runtime of traditional optimization methods such as EM soars. To learn the model, we used all 15, 051 Pfam MSAs from Yang *et al*. [20] subsampled down to 1,024 sequences each; each MSA is associated with structure data which we used to determine contacting sites. We estimated phylogenetic trees using FastTree [21] and then applied our CherryML method to estimate *Q*_400_. Tree reconstruction was parallelized and took approximately 1 minute per tree. The CherryML method took only 14 minutes using 8 CPU cores: 2 minutes for counting transitions and 12 minutes for the PyTorch optimizer. Extrapolating the results from Figure 1b, as well as based on our theoretical complexity results (see Methods for the details), we estimated that using EM for this task would have taken approximately 100 CPU-years – a million times slower!

Before analyzing the properties of our estimated rate matrix *Q*_400_, we used simulations to determine whether our method had produced a reliable estimate. To this end, we simulated data using *Q*_400_ for all 15,051 families, and then applied our CherryML method to re-estimate *Q*_400_. Our method was able to produce accurate estimates of most entries. Transitions with just one substitution proved the easiest to recover (Figure 2a) – as expected, since they have higher rates – with a median error of 0.8%. Transitions with two substitutions were harder to estimate owing to their small rates, but were still well estimated, particularly those with a higher rate, exhibiting a median error of 20.3% (Figure 2b).

**Figure 2:**
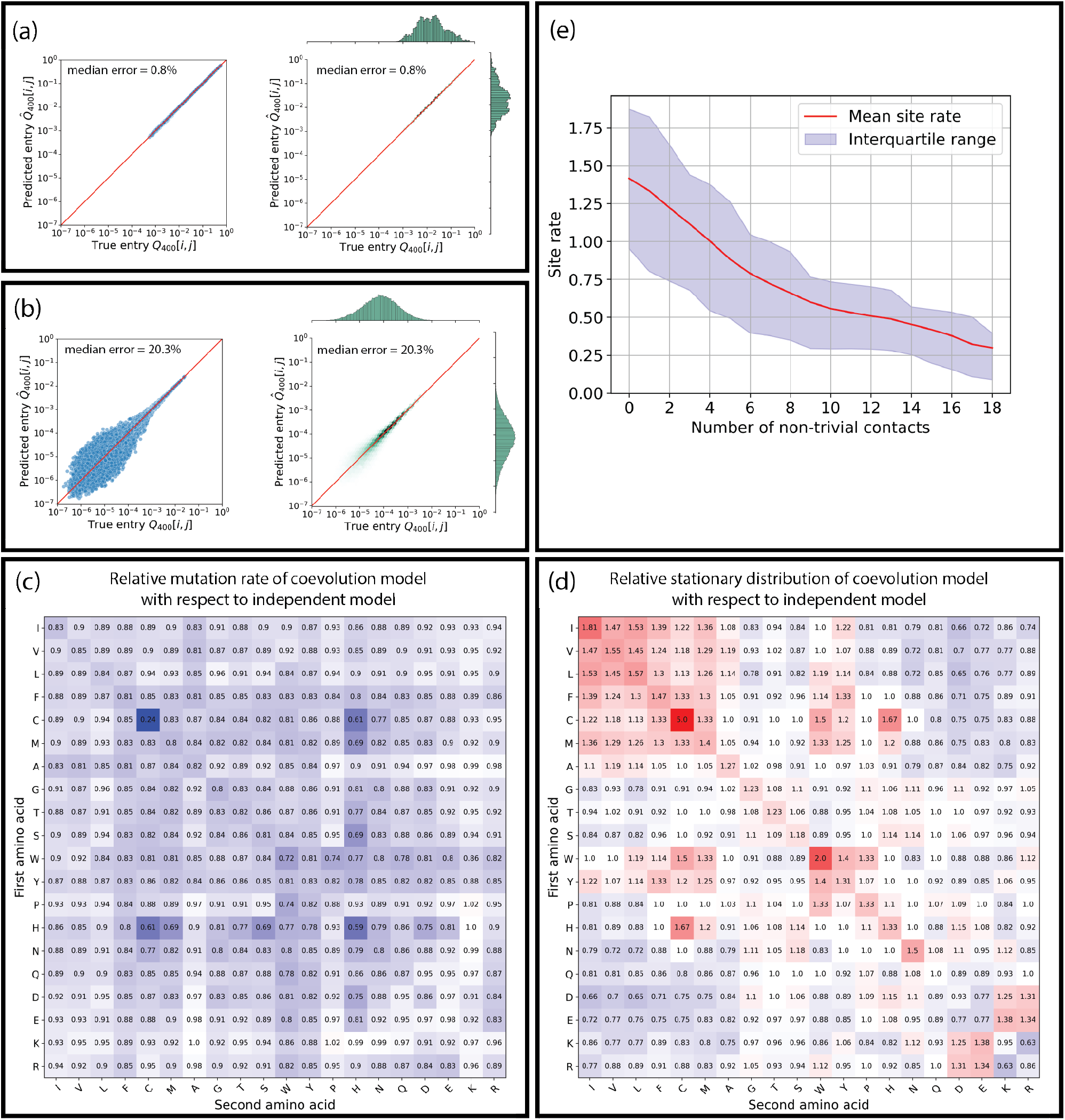
CherryML method applied to learn a 400 × 400 co-evolution model. Using simulated co-evolution data from 15,051 Pfam families each subsampled down to 1024 sequences, we verified that our method is able to accurately estimate co-transition rates for (a) single-site transitions (such as IL ↔ IA), and (b) the more challenging joint transitions (such as KE ↔ EK). The left plot is a scatterplot which reveals outliers, while the right plot is a density plot which shows that there are few outliers. (c) Mutation rates of co-evolution model differ from independent model, and recapitulate known biology such as the importance of disulfide bonds by assigning a significantly lower mutation rate to CC pairs. Residues are ordered by the hydropathy index. (d) Stationary distribution of co-evolution model differs from independent model, and recapitulates favorable residue pairings, such as hydrophobic amino acid pairs. (e) The more contacts a site has, the lower its mutation rate as estimated by FastTree [21].

We analyzed our estimated co-evolution model *Q*_400_ for amino acid pairs in contact and compared it with an independent model of amino acid evolution. We found that our model was able to learn the importance of disulfide bonds by assigning a low mutation rate to CC pairs, approximately 4 times smaller than what is predicted by an independent model (Figure 2c). Similarly, we found that our co-evolution model was able to learn the stability of certain kinds of residue contacts based on biochemical properties such as charge: our model assigns a higher observation probability to hydrophobic pairs (Figure 2d). The mutation rates and stationary distributions of our co-evolution model and the independent model are provided in Supplementary Figure S1 and Supplementary Figure S2. Our coevolution model has a global mutation rate of 1.51, which is much lower than the 2.0 predicted by a model of independent evolution. A retrospective analysis revealed that sites with more contacts have a lower mutation rate as estimated by FastTree. The trend is remarkably strong, as shown in Figure 2e. Finally, our model inferred interesting simultaneous substitutions at both sites in contact. For example, we observed that the KE ↔ EK transition probability is substantially higher in the co-evolution model than that in the independent model. This coupled evolution seems mediated by stability of electrostatic interaction and we found support for it in the original MSAs: for example, in the family `4kv7_1_A`, when sites 165 and 181 contain E and K, 97% of the time they are either KE or EK, while only 3% of the time they are EE or KK.

We envision that our CherryML framework will be broadly applicable to enable and scale up MLE under many models of molecular evolution. The ability to estimate transition rate matrices at unprecedented speed will transform the way that phylogenetic analysis is performed. Researchers will be able to estimate context-dependent rate matrices in a matter of minutes, such as by protein function, family, domain, or structure, which can then be applied – in place of the generic LG matrix – to estimate more accurate phylogenetic trees [9, 22]. In particular, recent advances in protein modeling, such as AlphaFold [23], have made such contextual structural information readily available.

Finally, although the CherryML method is most effective for reversible models of evolution – where it completely solves the problem of marginalizing out ancestral states – it can also be applied to irreversible models. In this case, the composite likelihood depends on the two branch lengths *t*_1_, *t*_2_ of each cherry, which can be quantized separately. For a typical model like LG [5], the terms in the quantized composite likelihood group together by pairs of quantized branch lengths (*τ*_*k*_, *τ*_*k*_′). As a result, EM can be performed in its classical form, except that now only *finitely* many ancestral state inferences need to be performed, thereby dramatically speeding up EM.

## Acknowledgments

Sebastian Prillo would like to acknowledge Alfredo Umfurer for many helpful discussions on software design. We would also like to thank Ian Holmes, John Huelsenbeck and Neil Thomas for helpful discussions. This research is supported in part by an NIH grant R35-GM134922.

## Methods

### Composite likelihood over cherries

The first idea of the CherryML method is to use a composite likelihood over cherries instead of the full likelihood. Let *D* = (*D*_1_, *D*_2_, …, *D*_*m*_) be the multiple sequence alignments and *T* = (*T*_1_, *T*_2_, …, *T*_*m*_) their associated phylogenetic trees. Then the log-likelihood of the data can be written as:

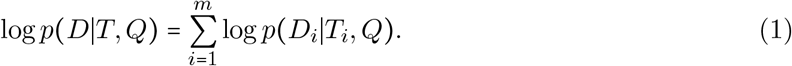

To obtain a composite likelihood, we select cherries 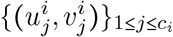 from each phylogenetic tree *T*_*i*_, where *c*_*i*_ is the number of cherries of *T*_*i*_, and then substitute the likelihood from Eq. (1) with the composite likelihood over the selected cherries:

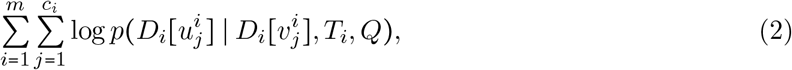

where *D*_*i*_ [*w*] is the row of *D*_*i*_ corresponding to leaf *w*. Cherries are considered in both directions, meaning that if *a, b* is a cherry then so is (*b, a*) (hence *c*_*i*_ is always even). We note that in the case of a reversible model, the likelihood term 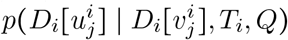 only depends on the distance between 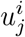 and 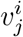 in the tree *T*_*i*_, so if we denote this distance as 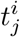, we can write, abusing notation slightly, the log-likelihood as:

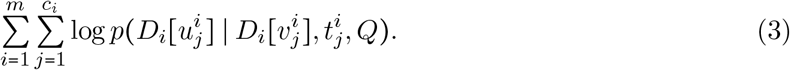

### Quantization of time

The second idea of the CherryML method is to quantize transition times. Concretely, we approximate each transition time *t* by one of the finitely many values *τ*_1_ < *τ*_2_ < ⋯ < *τ*_*b*_. We choose them to be geometrically spaced and to cover the whole operational range of transition times. As shown in Figure 1d, around 100 quantization points are enough to obtain negligible quantization error. We use *b* = 129 quantization points with a center value of *τ*_65_ = 0.03 and a geometric spacing of 1.1. Thus *τ*_1_ ≈ 6.7 × 10^−5^ and *τ*_129_ ≈ 13.4. This represents a wide range of transition times. We denote the quantized value of *t* as *q*(*t*), which is chosen to minimize the relative error, that is to say:

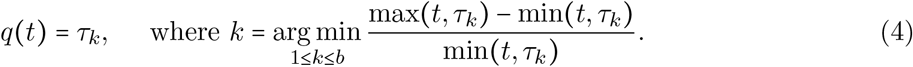

Quantization of a transition time *t* can be performed in time 𝒪(log *b*) with binary search. Any transition showcasing a transition time falling outside of the interval [*τ*_1_, *τ*_129_] (because it is either too small or too large) is discarded and thus dropped from the likelihood Eq. (3). Using a geometric spacing of 1.1 means that the relative error between successive points is 10%, and thus the *worst* quantization error (which happens in between two quantization points) is roughly 5%. The *average* quantization error is therefore around 2.5%.

In the case of a time-reversible model, quantizing time means replacing 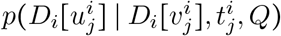 by 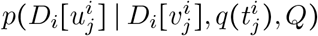. Concretely, we replace Eq. (3) with:

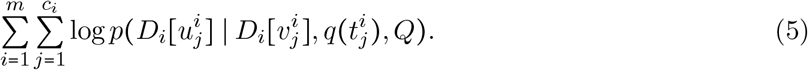

When this is done, the terms in the composite likelihood typically group together by quantized time *τ*_*k*_, and the cost of evaluating the likelihood no longer depends on the dataset size. We call this the *quantized (log-)likelihood*. For a model with site rate variation such as LG, transition times are adjusted by the site rate before quantizing.

### CherryML for the LG model

The LG model [5] of protein evolution posits that sites in a protein evolve independently under a scalar multiple of a rate matrix *Q* of size 20 × 20. The scalar for each site might be different and is called the *site rate*. If *l*_*i*_ denotes the length of the protein in family *i*, and 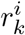 denotes the rate of site *k* (1 ≤ *k* ≤ *l*_*i*_), then time quantization under the reversible LG model is performed by replacing 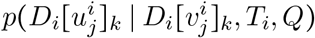 by 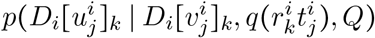. In other words, we obtain the quantized log-likelihood:

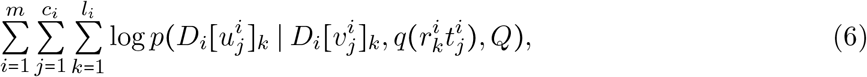

where *D*_*i*_[*w*]_*k*_ is site *k* at the row of *D*_*i*_ corresponding to leaf *w*. Transitions involving gaps are ignored. For the LG model, the terms of the quantized log-likelihood group together by quantized transition time *τ*_*k*_, and so Eq. (6) reduces to the simpler form:

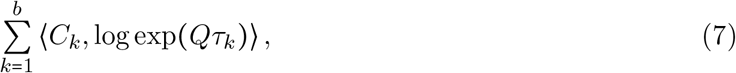

where *C*_*k*_ of size 20 × 20 is the frequency matrix for transitions between states occurring at a quantized transition time of *τ*_*k*_.

### CherryML for the co-evolution model

Our co-evolution model posits that a given pair of contacting sites in a protein sequence evolve together via a reversible 400 × 400 rate matrix *Q*. Since in a given protein a site might be in contact with more than one other site, we use a maximal matching to pair up contacting sites. This way, we obtain *p*_*i*_ pairs of sites 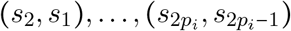 for family *i*. The quantized log-likelihood is thus:

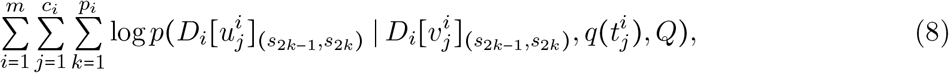

where 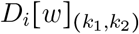 is the pair of site *k*_1_ and *k*_2_ at the row of *D*_*i*_ corresponding to leaf *w*. Because the order of the two sites does not matter (for example, the transition AL→ GI and the transition LA →IG are equivalent), when forming the composite likelihood we also augment the site pairs with the reverse pairs 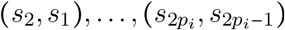. Transitions involving gaps are ignored. The log-likelihood Eq. (8) has the same functional form as LG model’s Eq. (7) except that *Q* is a 400 × 400 matrix instead of 20 × 20. Just like for the LG model, Eq. (8) reduces to the simpler form of Eq. (7) where *C*_*k*_ of size 400 × 400 is the frequency matrix for transitions between pairs of states occurring at a quantized transition time of *τ*_*k*_.

### Optimization with PyTorch

Optimization under the LG model and under the co-evolution model both require maximizing a log-likelihood of the form Eq. (7) with respect to a reversible rate matrix *Q* of size *s* × *s*. We achieve this using first-order optimization in PyTorch [18]. The reversible rate matrix *Q* is parameterized by a vector *θ* ∈ R^*s* × 1^ and an upper triangular matrix Θ ∈ R^*s* × *s*^, as follows: letting *π* = SoftMax(*θ*) and *S* = SoftPlus(Θ + Θ^⊺^), we take the off-diagonal entries of *Q* to be 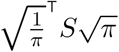 (where operations are performed entry-wise). The diagonal entries of *Q* are then uniquely determined. With this parameterization, optimization of *Q* translates to unconstrained optimization of *θ* and Θ. To ensure that the loss is on the same scale regardless of the dataset size, we divide the log-likelihood in Eq. (7) by the total number of transitions, that is to say the sum of all the entries in all the *C*_*k*_ matrices. For this we use the Adam optimizer [24] with a learning rate of 0.1. We train the single-site model for *g* = 2000 epochs and the co-evolution model for *g* = 500 epochs, and take the iteration with the lowest training loss as the final estimate. The matrix exponential layer is part of the PyTorch API and is based on an optimized Taylor approximation [25]. When benchmarking the performance of CherryML and EM as seen in Figure 1b,c, we use 1 CPU core for CherryML to make sure of a fair comparison of runtime between methods. Otherwise, we use 2 CPU cores to train the single-site model and 8 CPU cores to train the co-evolution model. We found these to be the optimal number of cores in our architecture; more CPU cores lead to communication overhead and overall slowdown. Initialization of the optimizer is described in the next section.

As a final remark, note that optimization of Eq. (7) with respect to an irreversible model is just as easy, by instead parameterizing *S* as *S* = SoftPlus(Θ_1_ + Θ_2_) where Θ_1_, Θ_2_ ∈ R^*s* × *s*^ are upper and lower triangular matrices respectively.

### Initialization with JTT-IPW

We initialize *θ* and Θ using a novel variant of the JTT method [2] that takes into account branch lengths, which we call *JTT with inverse propensity weighting*, or JTT-IPW for short. We observed that using the JTT-IPW initialization sped up convergence by up to an order of magnitude with respect to random initialization, and we used it when benchmarking both CherryML and EM. JTT-IPW is based on the Taylor expansion exp(*tQ*) = *I* + *tQ* + *o*(*t*) and on the decomposition *Q* = diag(*μ*) CTP where *μ*(*μ*_1_, …, *μ*_*s*_) is the vector of *mutabilities* and CTP is the matrix of *conditional transition probabilities*(except that the diagonal is filled with − 1). Here diag(*μ*) is the diagonal matrix with the entries of *μ* in the diagonal.

The JTT-IPW estimator of *Q* given the frequency matrices (*C*_1_, …, *C*_*b*_) and the transition times (*τ*_1_, …, *τ*_*b*_) is constructed as follows. First, to ensure that all quantities below are well defined, we add a small number of pseudocounts *ϵ* to the data entry-wise: *C*_*k*_ ← *C*_*k*_ + *ϵ* where *ϵ* = 10^−8^. Next, to ensure a reversible estimate of *Q* regardless of what the count matrices are, we symmetrize the count matrices via 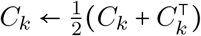. Now let 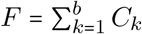 be the total transition frequency matrix. We first estimate the conditional transition probability matrix 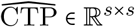 as follows:

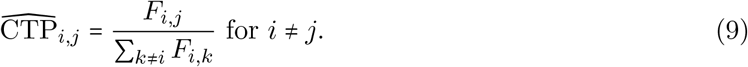

We set 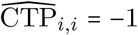 such that each row of 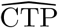 adds up to zero, as in a rate matrix. Finally, we estimate the mutability 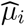 of each state using inverse propensity weighting as follows:

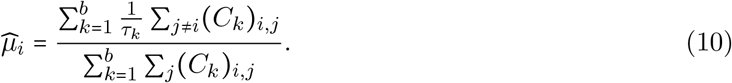

Finally, the JTT-IPW estimator of *Q* is given by:

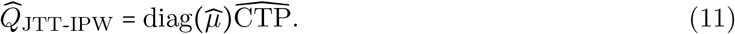

One can verify that 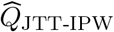 is a reversible rate matrix by noting that since *F*_*i,j*_ = *F*_*i,j*_, then 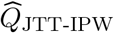 satisfies the detailed balance equation 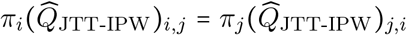 for all states *i, j*, where

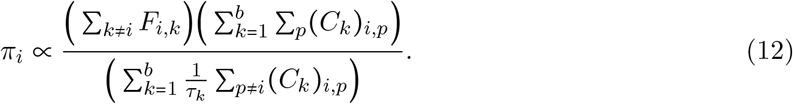

The reversibility of 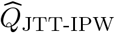 and the fact that all off-diagonal entries of 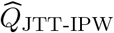 are positive (thanks to the pseudocounts) ensure that *θ*, Θ can be solved for. Concretely, we first solve for the stationary distribution *π* of 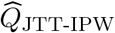 via an SVD decomposition. Then, we compute 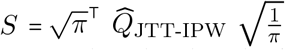. From *π* and *S* we can compute *θ* and Θ by taking *θ*_*i*_ = log(*π*_*i*_) and Θ_*i,j*_ = log(exp(*S*_*i,j*_) − 1) 𝟙{*i*< *j* }.

As for the intuition of why JTT-IPW works well as an initialization: as dataset size increases (in a suitable manner), one can show that 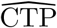 converges to the true conditional probability transition matrix. The mutabilities 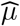 exhibit bias but it tends to 0 as branch lengths tend to 0.

### Computing the count matrices *C*_*k*_

The computational bottleneck of the CherryML optimizer is typically computing the count matrices *C*_*k*_. However, this is a simple and embarrassingly parallel task, for which we wrote a fast distributed-memory C++ implementation using MPI. To compute the transition frequency matrices *C*_*k*_ from the *m* multiple sequence alignments *D*_*i*_ and trees *T*_*i*_ using *p* MPI ranks, we give each worker *i* approximately *m*/*p* families for which it computes the count matrices 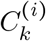. Once all workers are done, we compute 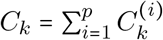. Time quantization is performed with binary search, introducing the log *b* factor in the 𝒪(*mnl* log *b*) time complexity.

When benchmarking the performance of CherryML and EM as seen in Figure 1b,c, we used 1 CPU core to count transitions for CherryML. Otherwise, we used 8 CPU cores.

### FastTree

We used FastTree [21] to estimate trees. FastTree was compiled to use double-precision arithmetic with:

~~~
gcc -DNO_SSE -DUSE_DOUBLE -O3 -finline-functions -funroll-loops
~~~

The number of rate categories was set using the `-cat` argument.

### Historian

We used the EM implementation provided by the Historian package [19]. We ran Historian’s `fit` command with the following arguments:

~~~
-band 0 -fixgaprates -mininc 0.000001 -maxiter 100000000 -nolaplace
~~~

The explanation for these arguments is as follows. Since the alignment is already provided, we set `-band 0`. Since there are no gaps in the data and we do not want to model gaps, we set `-fixgaprates`, and in the model file we set `insrate, delrate, insextprob, delextprob` all to zero. We set `-mininc 0.000001` to ensure that EM converges to a good solution. To avoid early termination due to reaching a maximum number of steps, we set `-maxiter 100000000`. Since we do not use pseudocounts during optimization, we set `-nolaplace`. Historian was run with ground truth trees specified in the Stockholm file format with the `#=GF NH` identifier. Site rate variation was handled by first splitting each MSA into one MSA per site rate, and scaling the tree accordingly by the site rate for each sub-MSA (this is the reduction described in the LG paper [5]).

### MSA Preprocessing

We preprocessed the MSAs in the Pfam dataset [20] by subsetting only sites that were aligned against residues in the reference sequence (which is the sequence for which we have structure data). In other words, we removed all sites aligned against gaps in the reference sequence. The length of the resulting MSA is therefore equal to the length of the reference sequence (which has no gaps).

### Protocol for Figure 1b,c

After preprocessing the MSAs from [20] as described in MSA Preprocessing, among all 15,051 families, we selected those with proteins having length between 190 and 230 sites inclusive, and at least 128 sequences. We then subsampled each MSA down to 128 sequences uniformly at random, making sure to sample the reference sequence. We then ran FastTree [21] with the LG matrix and to estimate trees for each family. We did not use site rate variation. Having estimated these realistic-looking trees, we then proceeded to simulate MSAs for each family. To do this, we simulated each site independently under the LG matrix. The root state was sampled from the stationary distribution of the LG matrix. Having simulated MSAs for all families, we then selected a random increasing sequence of family sets ℱ_4_ ⊂ ℱ_8_ ⊂ … ⊂ ℱ_1024_ which were used to train CherryML and EM. CherryML and EM were run with access to the ground truth trees. For CherryML, we formed the composite log-likelihood as described in CherryML for the co-evolution model and optimized for *Q* as described in Optimization with PyTorch. For EM, we used the Historian package as described in Historian. Our simulation scheme ensures that all MSAs are roughly of the same size, such that doubling the number of families is approximately equal to doubling the amount of Fisher information, which is important to obtain a reliable estimate of the relative efficiency of CherryML as compared to EM.

### Protocol for Figure 1d

After preprocessing the MSAs from [20] as described in MSA Preprocessing, for each of the 15,051 families, we subsampled them down to 1,024 sequences uniformly at random, making sure to sample the reference sequence. We then ran FastTree [21] with the LG matrix and 4 rate categories to estimate trees for each family as well as the site-specific rates. Having estimated these realistic-looking trees and site-specific rates, we then proceeded to simulate MSAs for each family. To do this, we simulated each site independently under the LG matrix and with the site rates estimated by FastTree. The root state was sampled from the stationary distribution of the LG matrix. CherryML was run with access to the ground truth trees and site rates. We formed the composite log-likelihood as described in CherryML for the co-evolution model and optimized for *Q* as described in Optimization with PyTorch. We explored varying the number of quantization points *b* while keeping the center value *τ*_⌈*b*/2⌉_ =0.03 and the range [*τ*_1_, *τ*_*b*_] approximately the same for all quantization points. For this, we chose the geometric increments [445.79, 21.11, 4.59, 2.14, 1.46, 1.21, 1.1, 1.048, 1.024] for quantization points 3, 5, 9, 17, 33, 65, 129, 257, 513 respectively.

### Protocol for Figure 1e

We followed the testing protocol described in the LG paper [5] to evaluate rate matrices. This means running PhyML [6] with the following arguments:

~~~
--nclasses 4 --datatype aa --pinv e --r_seed 0 --bootstrap 0 -f m \
--alpha e --print_site_lnl
~~~

The average improvement in per-site log-likelihood over JTT is shown. Thus, we evaluated the following four rate matrices: JTT [2], WAG [3], LG [5], and LG re-estimated using our CherryML method. To re-estimate LG using the CherryML method, we first used FastTree [21] with 4 rate categories as described in FastTree to estimate trees for each family as well as site-specific rates. The uninformative uniform rate matrix was used in FastTree. We then formed the composite log-likelihood as described in CherryML for the co-evolution model and optimized for *Q* as described in Optimization with PyTorch. The process was repeated, this time using the estimated *Q* to estimate trees with FastTree. This can be seen as coordinate ascent in tree and rate matrix space. The process was repeated 3 times until convergence, as typical.

### Determining contacting sites

To train our co-evolution model we used the Pfam dataset with structure data from [20]. A pair of sites was considered a non-trivial contact if (i) the distance between the beta carbons was less than 8 angstrom, and (ii) the distance in primary sequence was at least 7.

### Protocol for Figure 2c,d

After preprocessing the MSAs from [20] as described in MSA Preprocessing, for each of the 15,051 families, we subsampled them down to 1,024 sequences uniformly at random, making sure to sample the reference sequence. We then ran FastTree [21] with the LG matrix and 1 rate category to estimate trees for each family. We determined for each family which pairs of sites were in non-trivial contact as described in Determining contacting sites. We then used a maximal matching to pair up sites that were in contact. Finally, we formed the composite log-likelihood as described in CherryML for the co-evolution model and optimized for *Q* as described in Optimization with PyTorch. The independent model was obtained by training a single-site model on sites with at least one non-trivial contact, and then taking the product of the chain with itself.

### Protocol for Figure 2a,b

After preprocessing the MSAs from [20] as described in MSA Preprocessing, for each of the 15,051 families, we subsampled them down to 1,024 sequences uniformly at random, making sure to sample the reference sequence. We then ran FastTree [21] with the LG matrix and 1 rate category to estimate trees for each family. Having estimated these realistic-looking trees, we then proceeded to simulate MSAs for each family. To do this, we first determined for each family which pairs of sites were in non-trivial contact as described in Determining contacting sites. We then used a maximal matching to pair up sites that are in contact. Sites that are in contact were simulated under the co-evolution rate matrix *Q* estimated from real data as described in Protocol for Figure 2c,d. The root state was drawn from the stationary distribution of *Q*. CherryML was run with access to the ground truth trees.

### Protocol for Figure 2e

After preprocessing the MSAs from [20] as described in MSA Preprocessing, for each of the 15,051 families, we subsampled them down to 1,024 sequences uniformly at random, making sure to sample the reference sequence. We then ran FastTree [21] with the LG matrix and 20 rate categories to estimate trees for each family as well as the site-specific rates. We determined for each family which pairs of sites were in non-trivial contact as described in Determining contacting sites.

### Extrapolating the runtime of traditional methods for learning a co-evolution model

To estimate the time that it would take for traditional methods such as zeroth-order optimization or EM to estimate a 400 × 400 co-evolution rate matrix, we performed the following extrapolation. From Figure 1b we observe that it takes EM as implemented by Historian [19] around 30 CPU-hours to learn a single-site model on 1,024 families with 128 sequences each. Since the runtime of traditional methods scales linearly in the dataset size, this implies that learning a single-site model on all 15,051 families with approximately 1,024 sequences each would take on the order of 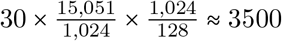 CPU-hours. However, runtime scales quadratically in the state space size, which means that increasing the state space size from *s* = 20 to *s* = 400 increases runtime by a factor of 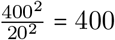. As a result, we estimate that learning a co-evolution model on all 15,051 families with approximately 1,024 sequences each would take on the order of 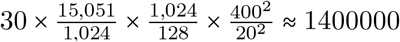 CPU-hours with traditional methods. The latter is approximately 160 CPU-years.

### Hardware Configuration

We used a node in Berkeley’s Savio cluster with 40 Intel Xeon Skylake 6230 @ 2.1 GHz cores and 384 GB RAM (which far exceeds our needs).

## Code Availability

Code for reproducing all results in this paper, as well as code implementing the CherryML method for the LG model and for the co-evolution model, will be made available on GitHub at the following repository:

https://github.com/songlab-cal/CherryML

## Data Availability

The LG paper [5] training and testing Pfam datasets consisting of 3,912 and 500 families respectively are available at:

http://www.atgc-montpellier.fr/models/index.php?model=lg

The Pfam dataset with structure data from [20] consisting of 15,051 families is located at:

https://files.ipd.uw.edu/pub/trRosetta/training_set.tar.gz

## Supplementary Figures

**Figure S1:**
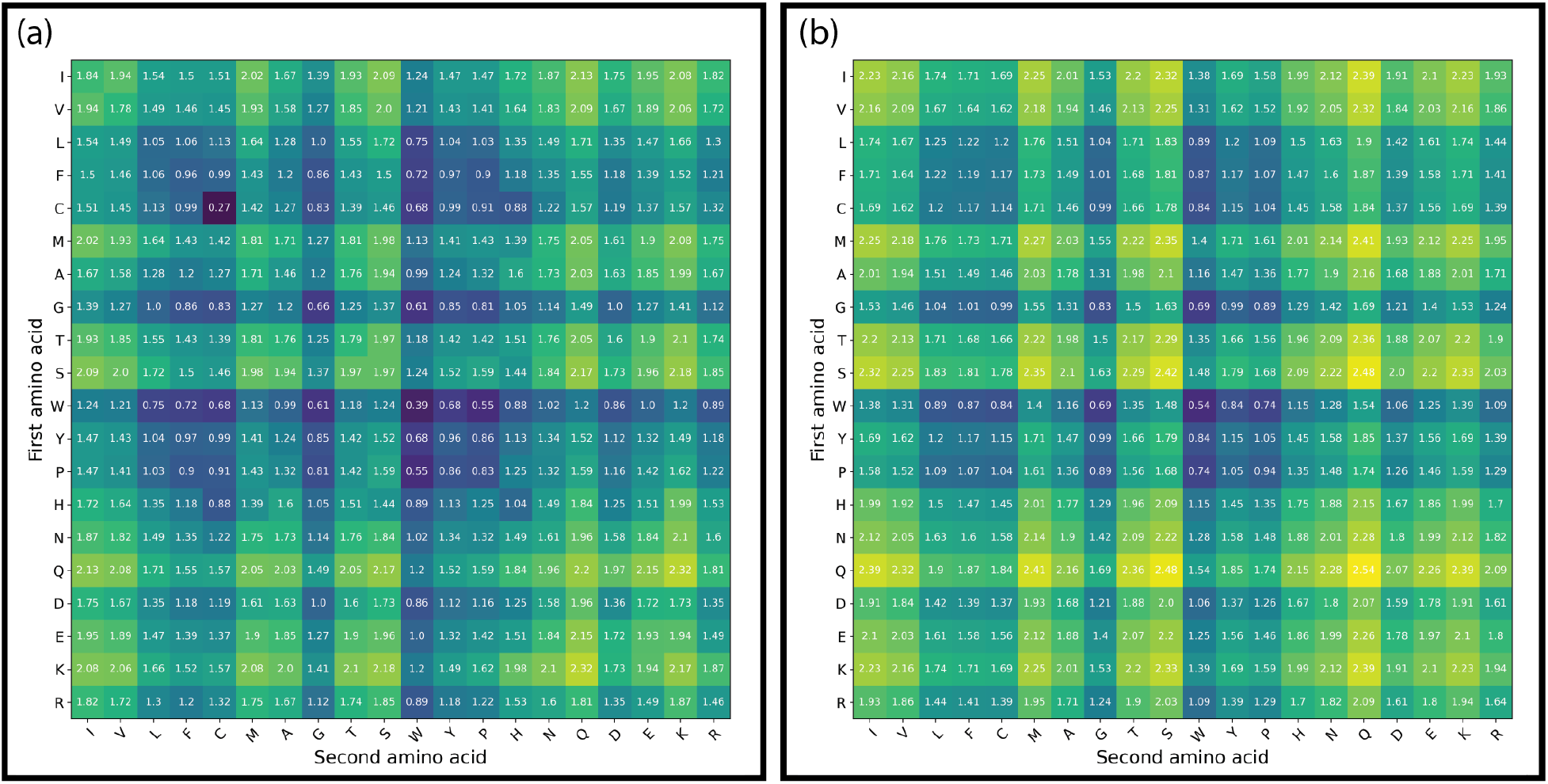
Mutation rates of (a) our 400 × 400 co-evolution model, and (b) the independent model.

**Figure S2:**
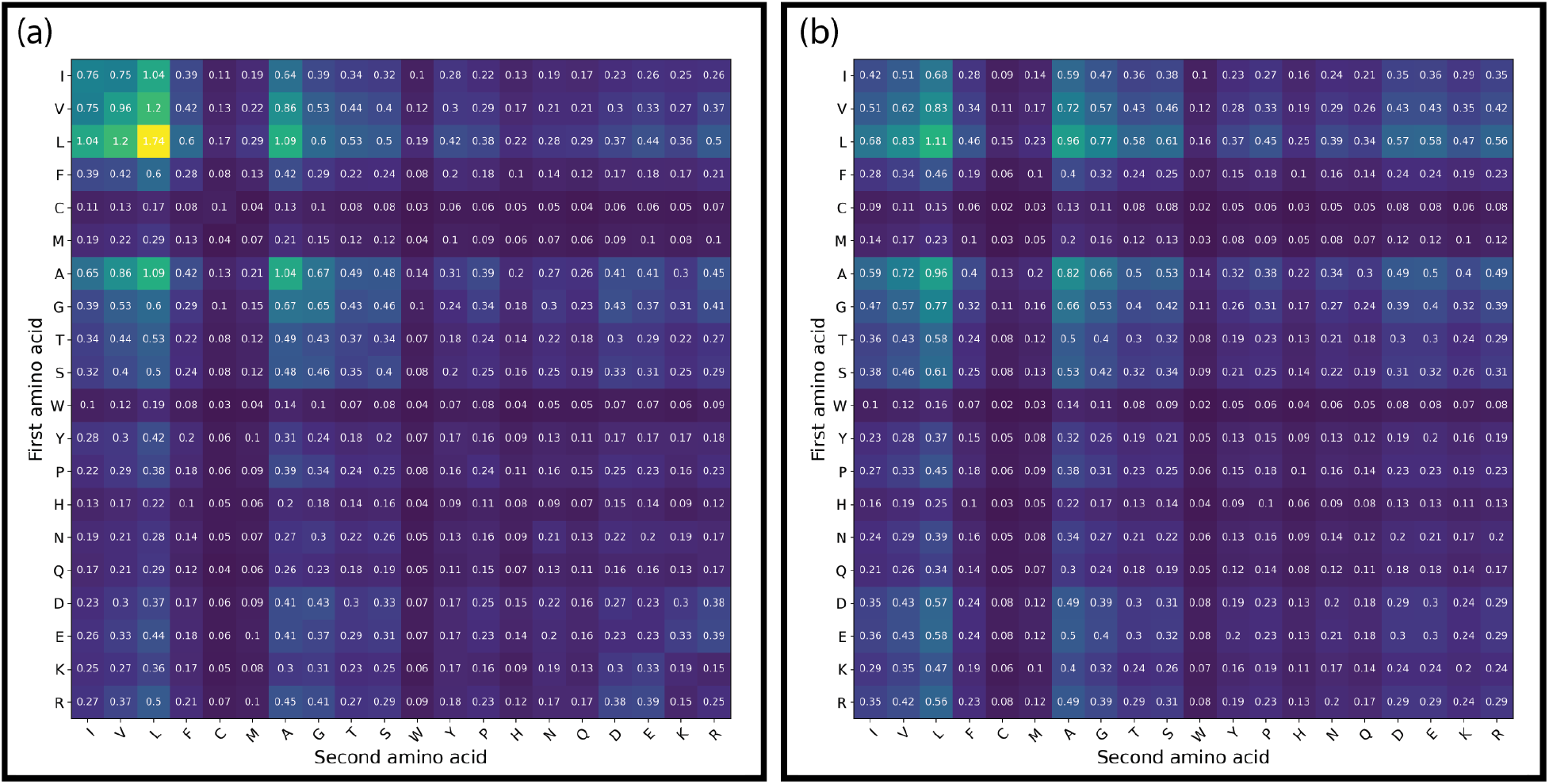
Stationary distribution of (a) our 400 × 400 co-evolution model, and (b) the independent model.

